## Supplementary material for "Quantitative definition of neurobehavior, vision, hearing and brain volumes in macaques congenitally exposed to Zika virus": S1 appendix

**S1 appendix.** Effect size and post-hoc sample size calculations SAS code.

```
*Read in raw data (csv format);
proc import file="data.csv" dbms=tab out=input;


data input;
set input;
proc sort;
by parameter group;

proc means noprint;
var value;
by parameter group;
output out=calcMean mean(value)=M SD(value)=SD N(value)=N;
run;


data control;
set calcMean;
if group="Control";
M1=M;
SD1=SD;
N1=N;
keep parameter  M1 SD1 N1;

data Zika;
set calcMean;
if group="Zika   ";
M2=M;
SD2=SD;
N2=N;
keep parameter M2 SD2 N2;

*Calculate Effect Sizes and conduct Post-Hoc Sample Size calculation;
data final;
merge control Zika;
by parameter
pooledSD=sqrt(((N1-1)*SD1**2+(N2-1)*SD2**2)/(N1+N2-1));

ES=abs(M2-M1)/pooledSD;

*Sample Size 80% power at two sided 0.05 significance level;
Ncalc=ceil(2*(((quantile("normal",0.8))
+quantile("normal",1-0.05/2))/ES)**2);


keep parameter ES Ncalc;

*Export estimated effect sizes;
proc export file="dataEffectSize.txt" replace;
run;
```
