## Supplementary figures and images for "Quantitative definition of neurobehavior, vision, hearing and brain volumes in macaques congenitally exposed to Zika virus"

### S1 figure

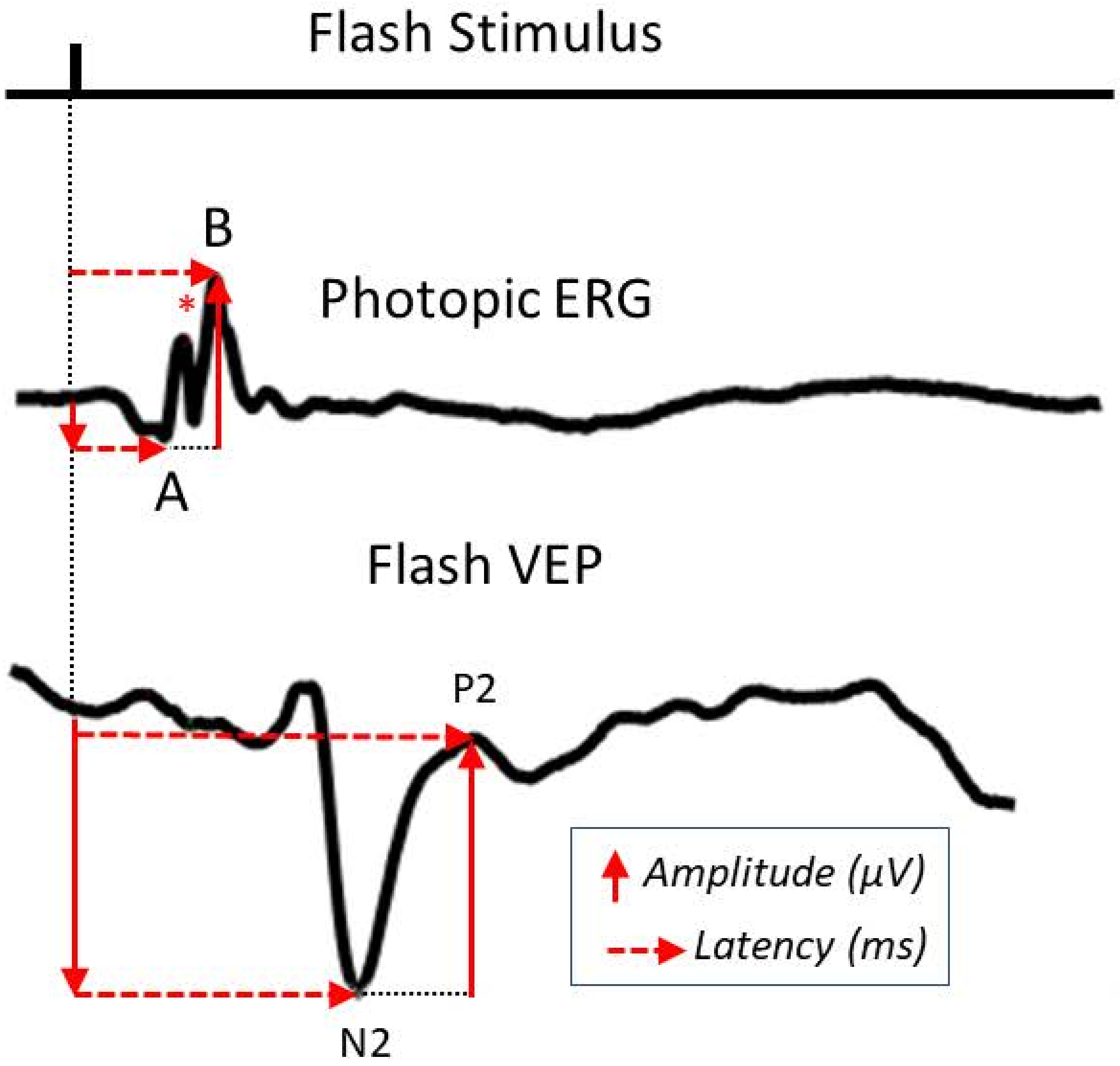

### S2 figure

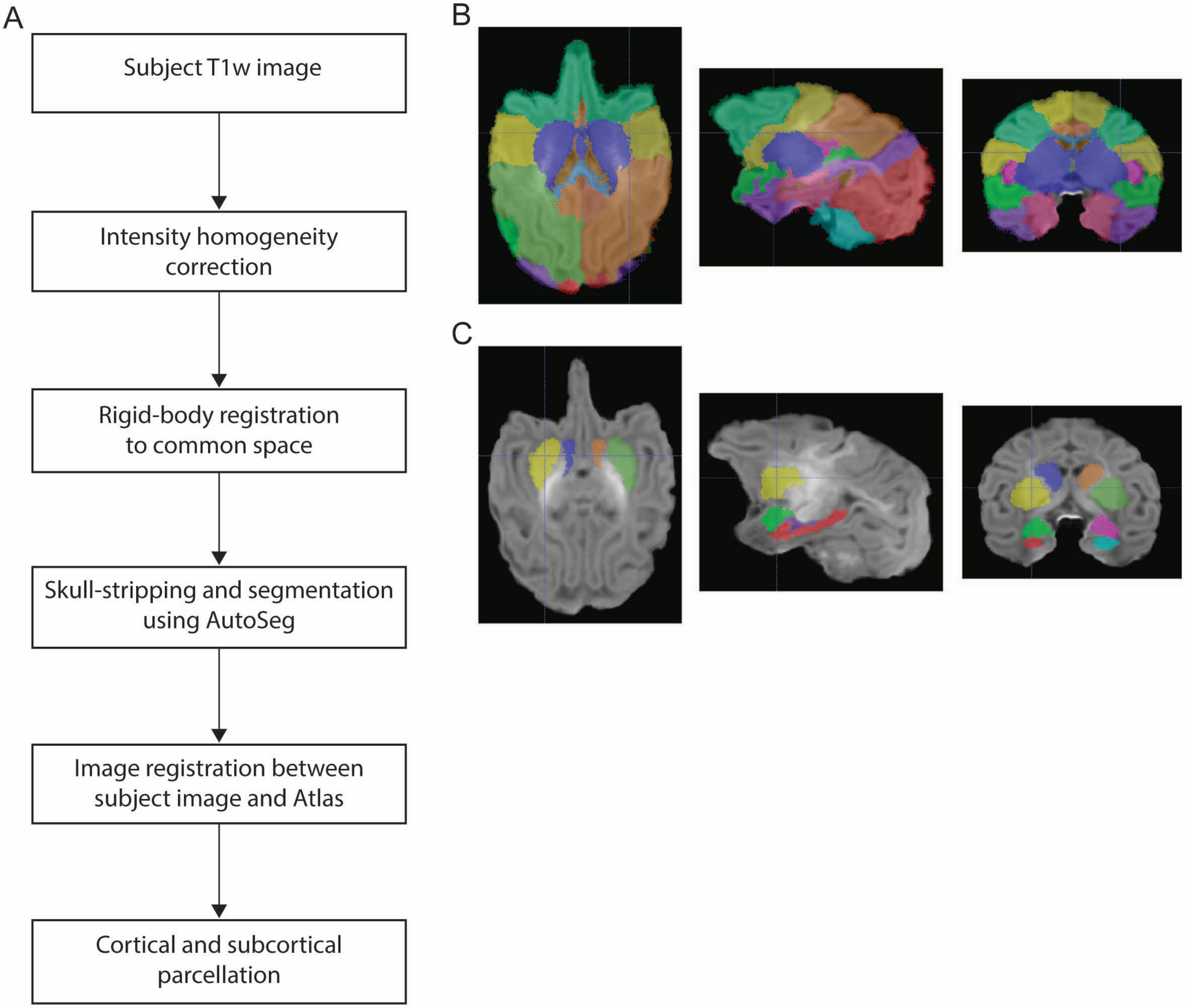

### S3 figure

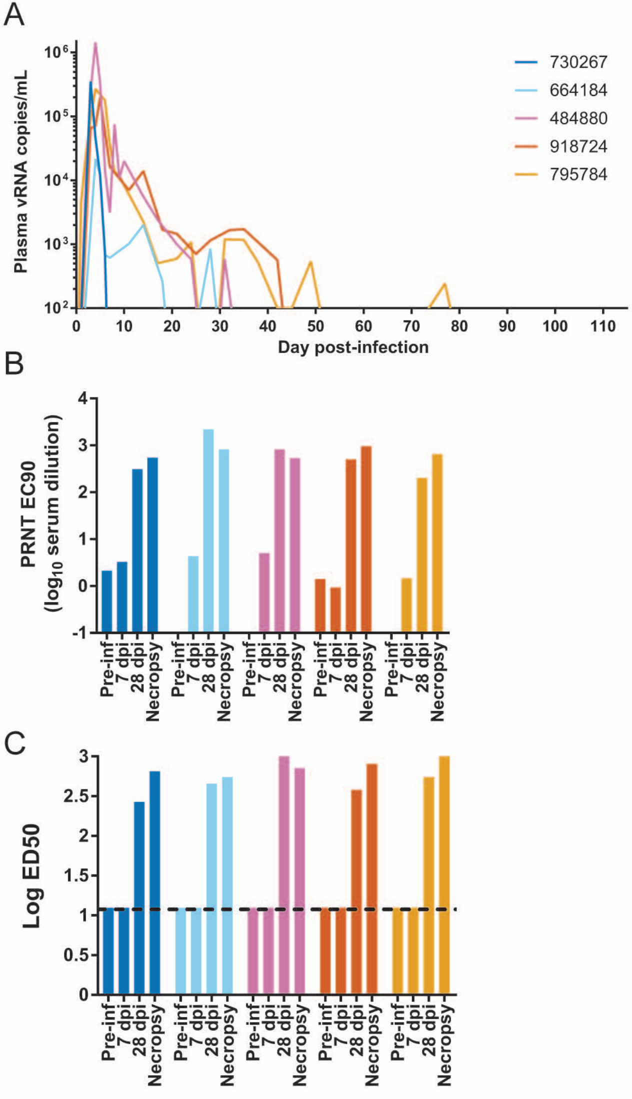

### S4 figure

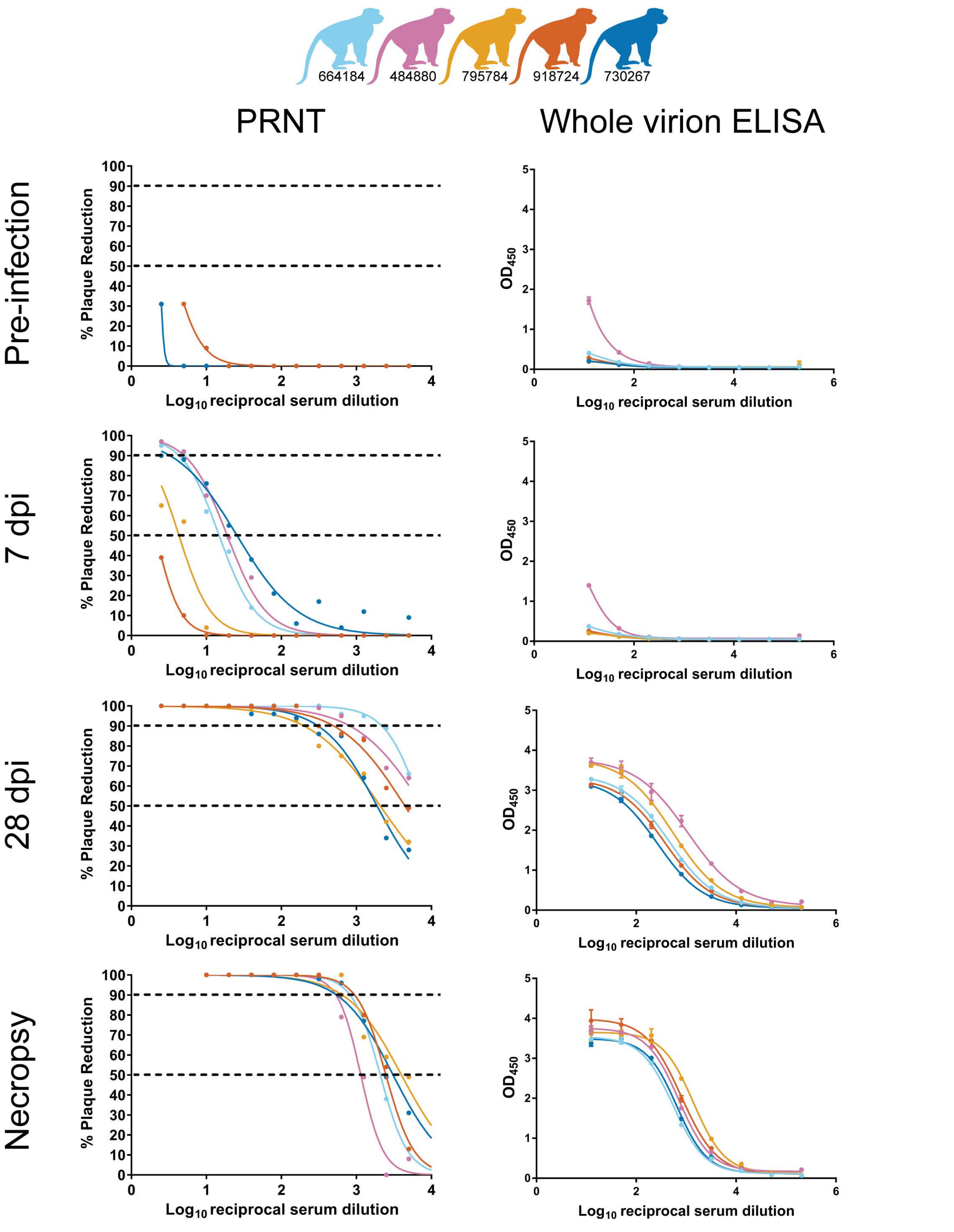

### S5 figure

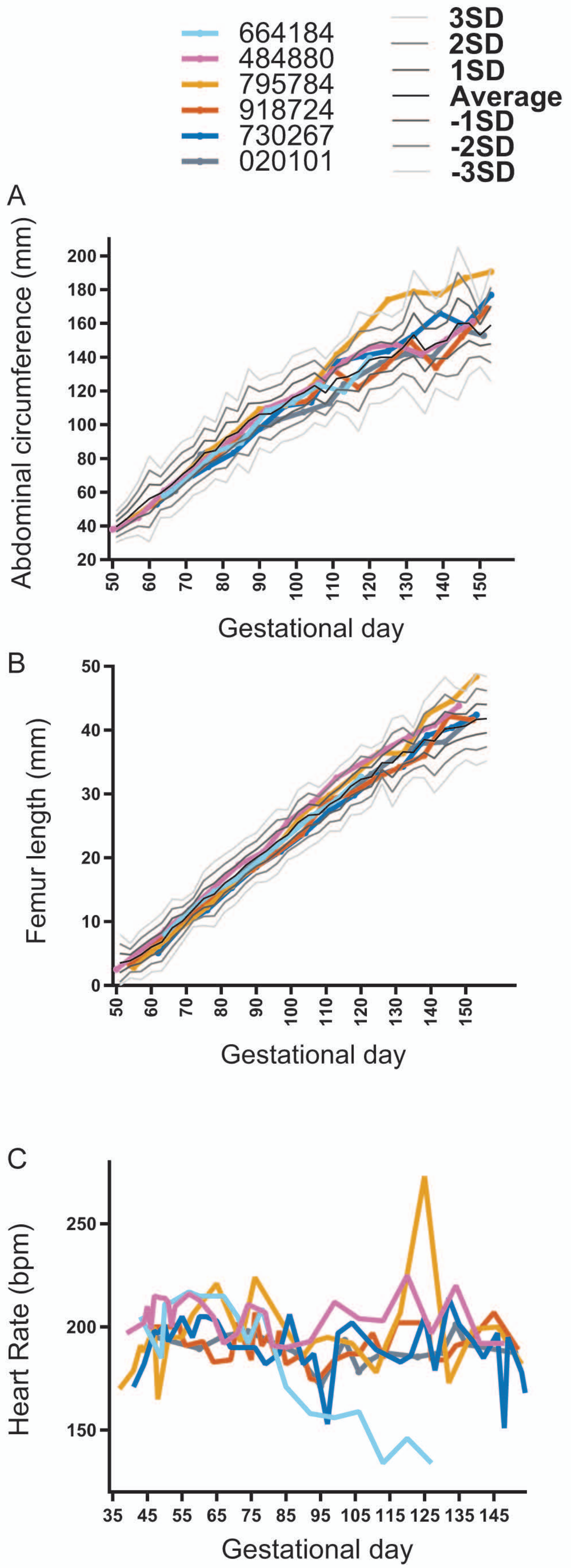

### S7 figure

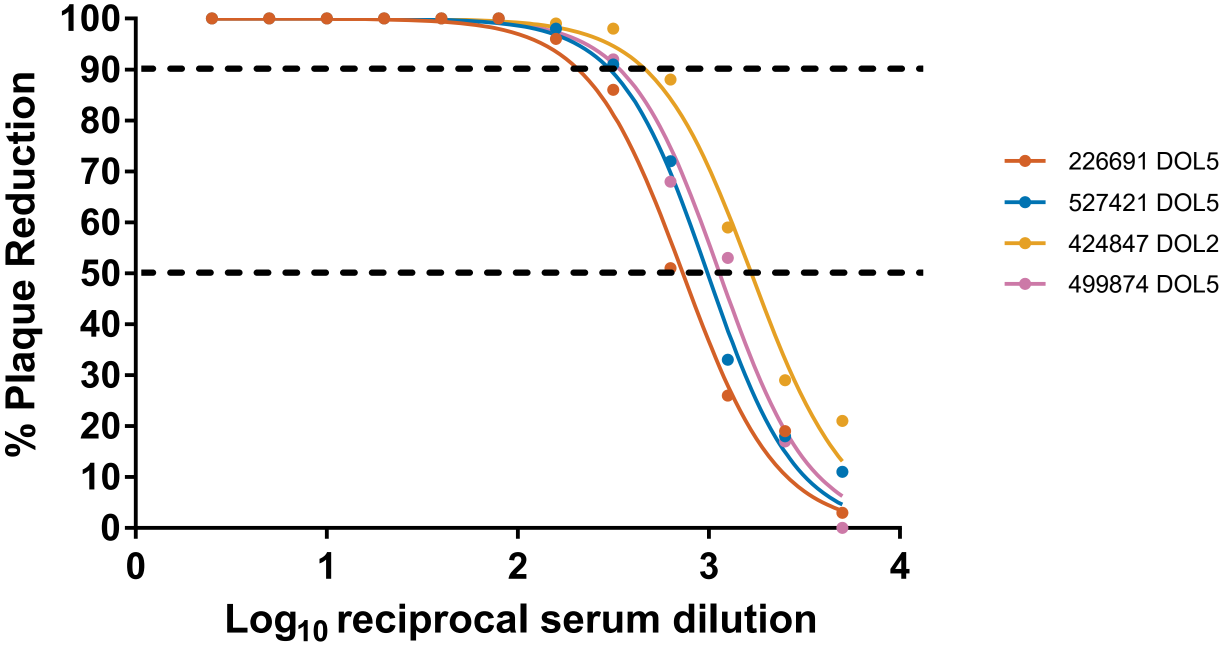

### S8 figure

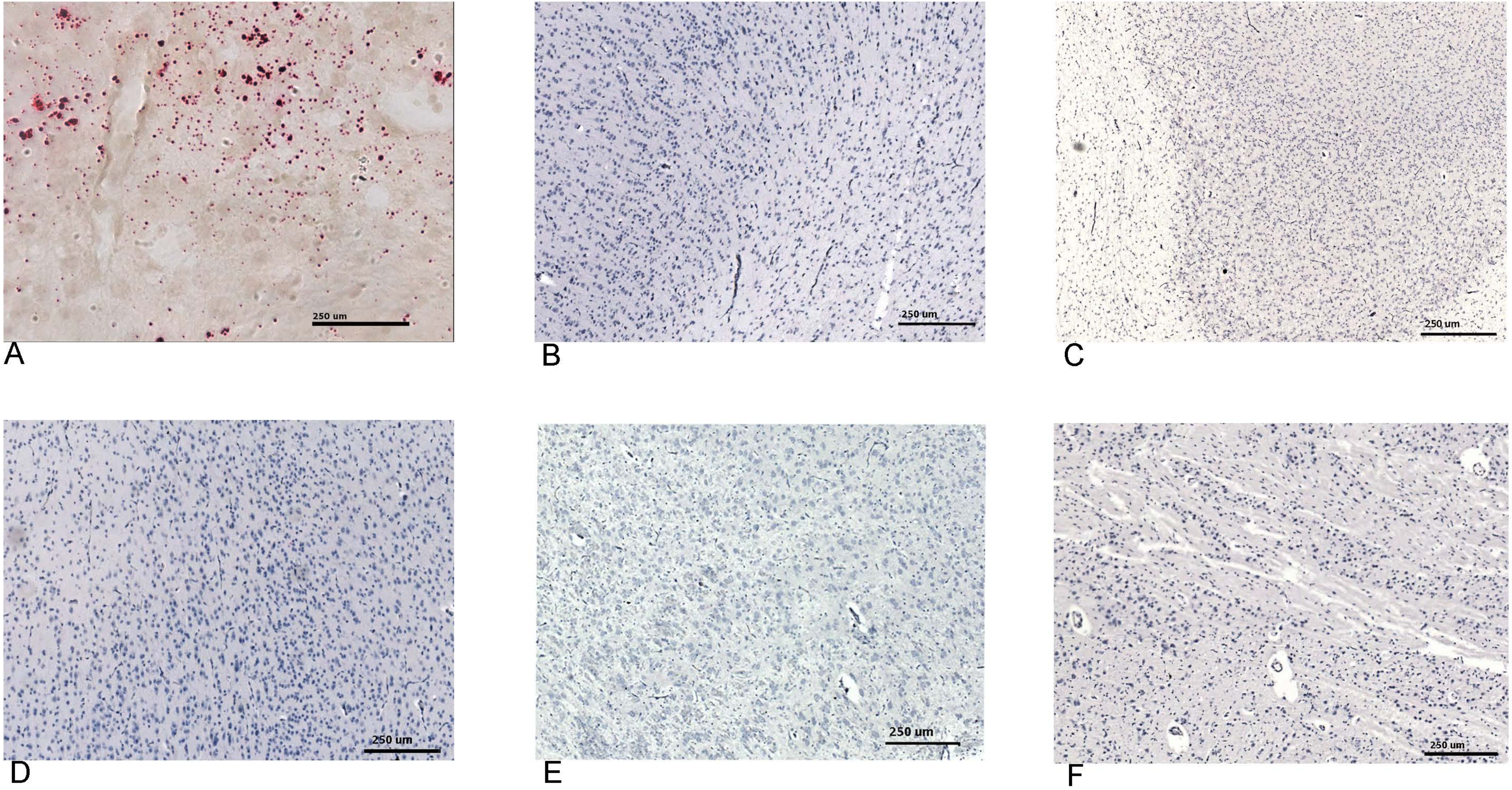
