## Supplementary material for "Quantitative definition of neurobehavior, vision, hearing and brain volumes in macaques congenitally exposed to Zika virus": S6 figure

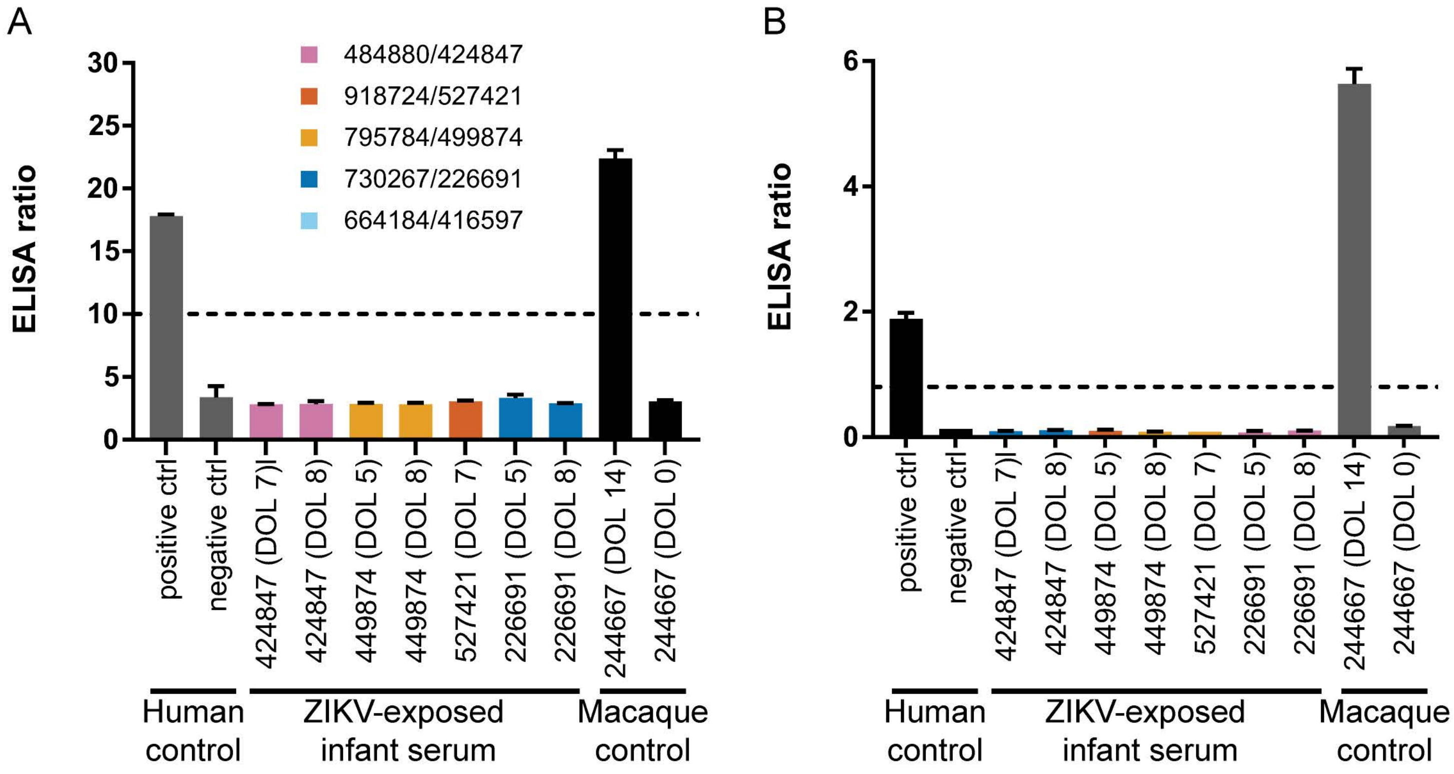


C

| Infant ages when negative viral loads were measured in body fluids (days of life) | | | | |
|---|---|---|---|---|
| Animal ID | Cord blood plasma | Plasma | Urine | CSF |
| 416597 | NA | NA | NA | NA |
| 424847 | 0 | 2, 8 | 1, 3, 7 | 8 |
| 499874 | 0 | 5, 7 | 1, 4, 5 | 7 |
| 527421 | 0 | 5, 8 | 1, 4, 6 | 8 |
| 226691 | 0 | 5, 8 | 1, 5, 6 | 8 |

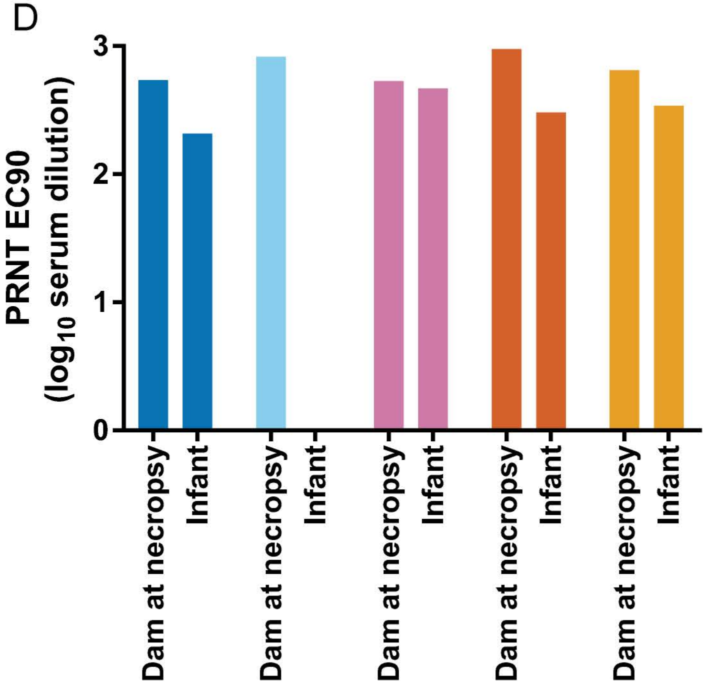
