## Supplementary material for "Quantitative definition of neurobehavior, vision, hearing and brain volumes in macaques congenitally exposed to Zika virus": S1 table

Supplementary Table 1. Infant ages at examination/evaluation and infant sex.

| Treatment Group | Infant ID | Examination | Gestational age at delivery (days) [165 days is term] | Postnatal age at exam (days of life) | Corrected gestational age at exam (gestational days) | Sex |
| --- | --- | --- | --- | --- | --- | --- |
| ZIKV-exposed | 416597 | Necropsy with histopathology | 133 | 133 | 133 | male |
|  | 424847 | Eye exam (OCT, visual electrophysiology, ophthalmic exam) | 155 | 2 | 157 | male |
|  |  | Hearing exam |  | ND | NA |  |
|  |  | Brain MRI |  | 7 | 162 |  |
|  |  | Neurobehavioral assessments |  | 1, 3, 6 | 156, 158, 161 |  |
|  |  | Necropsy with histopathology |  | 8 | 163 |  |
|  |  | Feeding volumes and weight gain |  | 0-8 | 155-163 |  |
|  | 499874 | Eye exam (OCT, visual electrophysiology, ophthalmic exam) | 155 | 5 | 160 | male |
|  |  | Hearing exam |  | ND | NA |  |
|  |  | Brain MRI |  | 7 | 162 |  |
|  |  | Neurobehavioral assessments |  | 1, 4, 6 | 156, 159, 161 |  |
|  |  | Necropsy with histopathology |  | 7 | 162 |  |
|  |  | Feeding volumes and weight gain |  | 0-7 | 155-162 |  |
|  | 527421 | Eye exam (OCT, visual electrophysiology, ophthalmic exam) | 154 | 5 | 159 | female |
|  |  | Hearing exam |  | ND | NA |  |
|  |  | Brain MRI |  | 7 | 161 |  |
|  |  | Neurobehavioral assessments |  | 1, 4, 6 | 155, 158, 160 |  |
|  |  | Necropsy with histopathology |  | 8 | 162 |  |
|  |  | Feeding volumes and weight gain |  | 0-8 | 154-162 |  |
|  | 226691 | Eye exam (OCT, visual electrophysiology, ophthalmic exam) | 155 | 5 | 160 | female |
|  |  | Hearing exam |  | 5 | 160 |  |
|  |  | Brain MRI |  | 7 | 162 |  |
|  |  | Neurobehavioral assessments |  | 1, 4, 6 | 156, 159, 161 |  |
|  |  | Necropsy with histopathology |  | 8 | 163 |  |
|  |  | Feeding volumes and weight gain |  | 0-8 | 155-163 |  |
| Control | 020501 | Eye exam (OCT, visual electrophysiology, ophthalmic exam) | 157 | 6 | 163 | female |
|  |  | Hearing exam |  | 2 | 159 |  |
|  |  | Brain MRI |  | 2 | 159 |  |
|  |  | Neurobehavioral assessments |  | 1, 5, 7 | 158, 162, 164 |  |
|  |  | Feeding volumes and weight gain |  | 0-8 | 157-165 |  |

Not done (ND). Not applicable (NA). Optical coherence tomography (OCT). Magnetic resonance imaging (MRI).
