## Supplementary material for "Quantitative definition of neurobehavior, vision, hearing and brain volumes in macaques congenitally exposed to Zika virus": S2 table

Supplementary Table 2. SNAP Constructs.

| SNAP Constructs | | | |
| --- | --- | --- | --- |
| Motor Maturity & Activity | Orientation | State Control | Sensory |
| Response Speed | Visual Orient- Right | Inversion | Tactile Response- RA |
| Motor Activity | Visual Orient- Left | Response Intensity | Tactile Response- LA |
| Coordinate | Visual Orient- Up | Soothability | Tactile Response- RL |
| Spontaneous Crawl | Visual Orient- Down | Tremulousness | Tactile Response- LL |
| Passive | Visual Follow- H | One Minute Vocalization Count Calculation | Galants- Right |
| Maintenance Balance | Visual Follow- V | Irritability | Galants- Left |
| Active Power | Duration of Looking | Consolability | One Min Voc Calculation |
|  | Attention | Struggle During Test | Calming Self |
|  | Orient to Auditory- R | Predominate State | Parachute |
|  | Orient to Auditory- L | Restrain | Rotation Test- Head Free |
|  | Distractible | Calming Self | Rotation Test- Head Held |
|  |  |  | Head Posture Prone |

Schneider Neonatal Assessment for Primates (SNAP). Right (R). Left (L). Horizontal (H). Vertical (V). Right arm (RA). Left arm (LA). Right leg (RL). Left leg (LL).
