## Supplementary material for "Quantitative definition of neurobehavior, vision, hearing and brain volumes in macaques congenitally exposed to Zika virus": S3 table

Supplementary Table 3. Infant procedural sedation medication record.

| Infant ID | Examination | Procedural sedation regimen |
| --- | --- | --- |
| 424847 | Brain MRI | Ketamine |
|  | Eye exam | ketamine, dexmedetomidine |
| 499874 | Brain MRI | ketamine, midazolam, propofol |
|  | Eye exam | midazolam, propofol, midazolam |
| 527421 | Brain MRI | propofol, isoflurane, midazolam, ketamine |
|  | Eye exam | propofol |
| 226691 | Brain MRI | isoflurane, midazolam, ketamine |
|  | Eye and hearing exam | isoflurane, ketamine, propofol, dexmedetomidine |
| 020501 | Brain MRI and hearing exam | dexmedetomidine, propofol |
|  | Eye exam | propofol, isoflurane, midazolam, ketamine |

Magnetic resonance imaging (MRI).
