## Supplementary material for "Quantitative definition of neurobehavior, vision, hearing and brain volumes in macaques congenitally exposed to Zika virus": S4 table

Supplementary Table 4. Histopathological description of decidua, placental bed and uterus, and placenta.

| Tissue Source | Organ System | Tissue Name | ZIKV-exposed | | | | | Control |
| --- | --- | --- | --- | --- | --- | --- | --- | --- |
|  |  |  | Dam: 664184 | Dam: 484880 | Dam: 795784 | Dam: 918724 | Dam: 730267 | Dam: 020101 |
|  |  |  | Fetus: 416597 | Infant: 424847 | Infant: 499874 | Infant: 527421 | Infant: 226691 | Infant: 020501 |
| Maternal | Reproductive | Decidua | Mild multifocal to diffuse neutrophilic deciduitis with multifocal necrosis. | Occasional rare aggregates of lymphocytes not diagnosed as deciduitis. | Mild multifocal lymphoplasmacytic to neutrophilic deciduitis. With Mild multifocal necrosis | Subacute to chronic lymphoplasmacytic deciduitis with multifocal necrosis and acute hemorrhages and mild-moderate, multifocal lymphoplasmacytic and neutrophilic vasculitis. | Moderate multifocal chronic lymphoplasmacytic and neutrophilic deciduitis. | Mild multifocal persistent muscularization of arteries and minimal multifocal chronic lymphoplasmacytic deciduitis. |
|  |  | Uterine placental bed | Focally extensive uterine diverticulum with severe segmental ischemic necrosis, severe diffuse necrosuppurative myometritis and endometritis, with perivascular hemorrhage, and vascular thrombosis | Minimal multifocal lymphocytic myometritis and mild multifocal random and perivascular lymphocytic endometritis. | Minimal lymphocytic endometritis. | There are no significant histologic lesions. | Moderate multifocal lymphoplasmacytic, perivascular myometritis. | There are no significant histologic lesions. |
| Fetal | Extraembryonic | Placenta | Multifocal severe ischemic necrosis of villi with mild multifocal to diffuse deciduitis. | Moderate multifocal regional basal plate infarction, acute coagulative necrosis, with neutrophilic villitis and intervillositis. | Multiple chronic (remote) infarctions with neutrophilic villitis and intervillositis. | Severe multifocal necrotizing and neutrophilic villitis and intervillositis with multifocal villous loss and collapse. | Moderate, multifocal decidual and basal plate persistent muscularization of arteries with multifocal fibrinoid necrosis, moderate lymphoplasmacytic and occasional neutrophilic vasculitis, moderate multifocal distal villous hypoplasia | Retroplacental hemorrhage, moderate multifocal basal thrombosis with ischemia and infarction with minimal multifocal intervillous fibrin and syncytial knots, focal avascular villi, mild multifocal subchorionic fibrin |
