## Supplementary material for "Quantitative definition of neurobehavior, vision, hearing and brain volumes in macaques congenitally exposed to Zika virus": S5 table

Supplementary Table 5. Infant clinical course summary.

|  | Fetus: 416597 | Infant: 424847 | Infant: 499874 | Infant: 527421 | Infant: 226691 | Infant: 020501 |
| --- | --- | --- | --- | --- | --- | --- |
| Resuscitation summary after delivery | Demise | Breathed spontaneously without support. | Poor respiratory effort at birth. Noninvasive positive pressure ventilation required to improve respiratory distress and increase SpO2 to >90% for first 60 minutes of life. Blow by oxygen sufficient to maintain SpO2 >90% after 1 hour of life. Blood glucose level 63 mg/dL shortly after birth. Administered furosemide x2 intramuscularly and 5% dextrose orally. Transferred to an oxygen cage in the nursery after stabilization. | Comfortable work of breathing at birth, but noted at 5 minutes of life to have SpO2 of 70-79%, so blow by oxygen was applied and SpO2 improved to >90%. She required intermittent blow by oxygen for the first 60 minutes of life to support to maintain SpO2 >90%. | No respiratory effort immediately after birth. After 5 breaths provided with a bag valve mask, she had spontaneous respiratory effort. She required intermittent blow by oxygen for the first 30 minutes of life to maintain Sp02 >90%. | Required transient blow by oxygen for low SpO2, which resolved by 10 minutes of life. |
| Assessment for congenital anomalies at birth (overlapping suture, redundant scalp skin, contractures) | Significant autolysis, unable to assess for congenital anomalies given condition | No congenital anomalies apparent on physical exam | No congenital anomalies apparent on physical exam | No congenital anomalies apparent on physical exam | No congenital anomalies apparent on physical exam | No congenital anomalies apparent on physical exam |
| Clinical course | NA | Housed in nursery. No respiratory or thermoregulation support needed | Housed in nursery. Required supplemental ambient oxygen for 20 hours after birth because of poor oxygenation and continued to require a heating pad in the incubator to maintain normothermia. Infant was feeding formula normally, was normothermic and had normal oxygen saturations prior to a sedated eye exam on DOL 5. Following this exam, infant required supplemental ambient oxygen for 4 hours for poor oxygenation and tachypnea. He was euthanized after his next sedated exam (a brain MRI) on day of life 7, earlier than was planned on day of life 8, because of the concern that there would be worsening respiratory distress. | Housed in nursery. No respiratory or thermoregulation support needed | Housed in nursery. No respiratory or thermoregulation support needed | Housed in nursery. No respiratory or thermoregulation support needed |
| Microbiology results | ND | ND | Blood cultures drawn on DOL 7 just prior to euthanasia were negative. Lung tissue sample at necropsy had a negative gram stain but grew moderate amounts of *Klebsiella oxytoca, Enterobacter cloacae, Staphylococcus aureus* after one day of culture. Cerebrospinal fluid, and liver, brain and spleen sections submitted for bacterial culture had no growth. | ND | ND | ND |
| Complete blood count | ND | ND | Obtained because of concern for infection on DOL 1 and 4. White blood cell count were within the normal range on DOL 1 and below the normal range on DOL 4. See Supplementary Table 5 for the specific values and additional CBC parameters. | ND | Obtained per routine measurement on DOL 7. See Supplementary Table 5 for specific CBC parameters. | ND |
| Clinically-indicated imaging | ND | ND | Chest radiograph and thoracic and abdominal ultrasounds were done on DOL 0 because of respiratory distress and had no abnormalities. | Echocardiogram was done on DOL 6 to follow up pericardial effusion seen in utero and had no abnormalities. | ND | ND |

Not done (ND). Not applicable (NA). Day of life (DOL). Complete blood count (CBC). Peripheral oxygen saturation (SpO_2_).
