## Supplementary material for "Quantitative definition of neurobehavior, vision, hearing and brain volumes in macaques congenitally exposed to Zika virus": S6 table

Supplementary Table 6. SNAP effect sizes and sample size estimates.

| Outcome | | Effect Size | Sample Size Allocation 1:1* | | | Sample Size Allocation 2:1* | | |
| --- | --- | --- | --- | --- | --- | --- | --- | --- |
|  |  |  | N: ZIKV | N: Ctr | Total N | N: ZIKV | N: Ctr | Total N |
| Motor Maturity & Activity | Day 1 | 1.56 | 8 | 8 | 16 | 12 | 6 | 18 |
|  | Day 3/4 | 5.57 | 2 | 2 | 4 | 4 | 2 | 6 |
|  | Day 6/7 | 0.8 | 26 | 26 | 52 | 39 | 20 | 59 |
| Orientation | Day 1 | 0.06 | 4362 | 4362 | 8724 | 6543 | 3272 | 9815 |
|  | Day 3/4 | 2.1 | 5 | 5 | 10 | 8 | 4 | 12 |
|  | Day 6/7 | 1 | 17 | 17 | 34 | 26 | 13 | 39 |
| Sensory | Day 1 | 1.18 | 13 | 13 | 26 | 19 | 10 | 29 |
|  | Day 3/4 | 0.17 | 545 | 545 | 1090 | 817 | 409 | 1226 |
|  | Day 6/7 | 1.2 | 12 | 12 | 24 | 18 | 9 | 27 |
| State Control | Day 1 | 0.61 | 44 | 44 | 88 | 65 | 33 | 98 |
|  | Day 3/4 | 0.17 | 545 | 545 | 1090 | 817 | 409 | 1226 |
|  | Day 6/7 | 0.48 | 70 | 70 | 140 | 104 | 52 | 156 |

Control (Ctr). * These sample size estimates were performed for the observed effect sizes with 80% power at the two-sided 0.05 significance level.
