## Supplementary material for "Quantitative definition of neurobehavior, vision, hearing and brain volumes in macaques congenitally exposed to Zika virus": S7 table

Supplementary Table 7. Ophthalmic exam summary of ZIKV-exposed and control infants.

| **Evaluation type** | **ZIKV-exposed infants** | | | | **Control infant** |
| --- | --- | --- | --- | --- | --- |
|  | **424847** | **499874** | **527421** | **226691** | **020501** |
| **Indirect ophthalmoscopy** |  |  |  |  |  |
| Pupillary response* | Slight response bilaterally | Unable to assess response adequately with anesthesia | Reactive | Reactive | Reactive |
| Eyelids and adnexa | Normal | Normal | Normal | Normal | Normal |
| Conjunctiva | White and quiet | White and quiet | White and quiet | White and quiet | White and quiet |
| Cornea | A subtle whorl-shaped corneal defect inferotemporally in the right cornea at the level of the anterior stroma. No corneal abrasion was present. The left cornea was normal. | Clear | Clear | Clear | Clear |
| Anterior chamber | Deep and quiet | Deep and quiet | Deep and quiet | Deep and quiet | Deep and quiet |
| Iris | Unremarkable | Unremarkable | Unremarkable | Unremarkable | Unremarkable |
| Lenses | Clear | Clear | Clear | Clear | Clear |
| Vitreous | No opacities | No opacities | No opacities | No opacities | No opacities |
| Hyaloid artery^ | Present bilaterally | Posterior hyaloid artery present bilaterally | Posterior hyaloid artery present bilaterally | Posterior hyaloid artery present bilaterally | Not described |
| Tunica vasculosa lentis | Not commented on | Persistent bilaterally | Persistent bilaterally | Persistent bilaterally | Not described |
| Optic nerve | Normal | Normal | Normal | Normal | Normal |
| Retina | No lesions | Subtle salt-and-pepper retinal pigmented epithelium mottling in both eyes, no large lesions | No lesions | No lesions | No lesions |
| Choroid | No lesions | No lesions | No lesions | No lesions | No lesions |
| **Intraocular pressure (mm Hg)#** |  | | | | |
| Right eye | 13 | 7 | 14 | 11 | 6 |
| Left eye | 14 | 8 | 14 | 10 | 7 |

Not applicable (NA). Not done (ND).

*Pupil exam is limited due to general anesthesia, so degree of reactivity cannot be determined, only that the pupils were reactive.

\# Normal ranges of neonatal macaque intraocular pressures are not known; only juvenile macaques (aged 7 months to 3 years) have been studied and they have an average intraocular pressure of 15.7 mm Hg (1).

^ The persistence of the tunica vasculosa lentis is a normal finding in newborn rhesus macaques (2) at corrected gestational ages close to the infants used in these ocular exams. A persistent hyaloid artery in newborn rhesus macaques is a normal finding and typically persists until the age of 2-3 weeks (3).
