## Supplementary material for "Quantitative definition of neurobehavior, vision, hearing and brain volumes in macaques congenitally exposed to Zika virus": S8 table

Supplementary Table 8. Ocular coherence tomography effect sizes and sample size estimates.

| Layer | Effect Size | Sample Size Allocation 1:1* | | | Sample Size Allocation 2:1* | | |
| --- | --- | --- | --- | --- | --- | --- | --- |
|  |  | N: ZIKV | N: Ctr | Total N | N: ZIKV | N: Ctr | Total N |
| Choroid | 0.39 | 105 | 105 | 210 | 157 | 79 | 236 |
| GCL | 1.64 | 7 | 7 | 14 | 11 | 6 | 17 |
| INL | 0.91 | 20 | 20 | 40 | 30 | 15 | 45 |
| IPL | 2.09 | 5 | 5 | 10 | 8 | 4 | 12 |
| ONL | 1.56 | 8 | 8 | 16 | 12 | 6 | 18 |
| OPL | 1.47 | 9 | 9 | 18 | 13 | 7 | 20 |
| Photoreceptor Inner Segment | 0.29 | 188 | 188 | 376 | 282 | 141 | 423 |
| Photoreceptor Outer Segment | 2.11 | 5 | 5 | 10 | 8 | 4 | 12 |
| RNFL | 0.74 | 30 | 30 | 60 | 45 | 23 | 68 |
| Total Retina | 0.36 | 123 | 123 | 246 | 184 | 92 | 276 |
| All layers combined | 1.16 | 13 | 13 | 26 | 20 | 10 | 30 |

Control (Ctr). * These sample size estimates were performed for the observed effect sizes with 80% power at the two-sided 0.05 significance level.
