## Supplementary material for "Quantitative definition of neurobehavior, vision, hearing and brain volumes in macaques congenitally exposed to Zika virus": S9 table

Supplementary Table 9. Visual electrophysiology effects sizes and sample sizes estimates.

| Component | Wave characteristic | Side | Effect Size | Sample Size Allocation 1:1* | | | Sample Size Allocation 2:1* | | |
| --- | --- | --- | --- | --- | --- | --- | --- | --- | --- |
|  |  |  |  | N: ZIKV | N: Ctr | Total N | N: ZIKV | N: Ctr | Total N |
| A wave | Amplitude | Left | 1.32 | 11 | 11 | 22 | 16 | 8 | 24 |
|  |  | Right | 1.55 | 8 | 8 | 16 | 12 | 6 | 18 |
|  | Latency | Left | 1.77 | 7 | 7 | 14 | 10 | 5 | 15 |
|  |  | Right | 0.01 | 156979 | 156979 | 313958 | 235468 | 117734 | 353202 |
| B wave | Amplitude | Left | 1.38 | 10 | 10 | 20 | 14 | 7 | 21 |
|  |  | Right | 0.57 | 50 | 50 | 100 | 74 | 37 | 111 |
|  | Latency | Left | 0.57 | 50 | 50 | 100 | 74 | 37 | 111 |
|  |  | Right | 0.14 | 802 | 802 | 1604 | 1203 | 602 | 1805 |
| N2 wave | Amplitude | Left | 1.14 | 14 | 14 | 28 | 20 | 10 | 30 |
|  |  | Right | 1.23 | 12 | 12 | 24 | 18 | 9 | 27 |
|  | Latency | Left | 0.13 | 930 | 930 | 1860 | 1395 | 698 | 2093 |
|  |  | Right | 0.89 | 21 | 21 | 42 | 32 | 16 | 48 |
| P2 wave | Amplitude | Left | 0.84 | 24 | 24 | 48 | 36 | 18 | 54 |
|  |  | Right | 0.83 | 24 | 24 | 48 | 36 | 18 | 54 |
|  | Latency | Left | 1.14 | 14 | 14 | 28 | 20 | 10 | 30 |
|  |  | Right | 1.18 | 13 | 13 | 26 | 19 | 10 | 29 |
| Root mean squared (RMS) | | Left | 0.64 | 40 | 40 | 80 | 60 | 30 | 90 |
|  |  | Right | 0.36 | 123 | 123 | 246 | 184 | 92 | 276 |
| All measurements combined | | | 0.87 | 22 | 22 | 44 | 33 | 17 | 50 |

Control (Ctr). * These sample size estimates were performed for the observed effect sizes with 80% power at the two-sided 0.05 significance level.
