## Supplementary material for "Quantitative definition of neurobehavior, vision, hearing and brain volumes in macaques congenitally exposed to Zika virus": S10 table

Supplementary Table 10. Cortical and subcortical brain region volumes with sample size estimates, corrected by total brain volume or intracranial volume.

| Brain regions | | Brain region volumes | | | | | Sample Size Estimates* | | | | | | |
| --- | --- | --- | --- | --- | --- | --- | --- | --- | --- | --- | --- | --- | --- |
|  |  |  |  |  |  |  | Observed Effect Size | Sample Size Allocation 1:1 | | | Sample Size Allocation 2:1 | | |
|  |  | 424847 | 499874 | 527421 | 226691 | 020501 |  | N: ZIKV | N: Ctr | Total N | N: ZIKV | N: Ctr | Total N |
| Entire brain regions (mm^3^) | Total brain volume | 49537.70 | 49137.00 | 49558.10 | 51717.20 | 53875.10 | 1.93 | 6 | 6 | 12 | 8 | 4 | 12 |
|  | Total CSF | 9491.64 | 8603.88 | 9455.10 | 10083.40 | 10908.80 | 1.76 | 7 | 7 | 14 | 10 | 5 | 15 |
|  | R Lat Ventricle | 47.90 | 101.96 | 88.45 | 72.53 | 104.63 | 1.15 | 13 | 13 | 26 | 20 | 10 | 30 |
|  | L Lat Ventricle | 80.35 | 83.34 | 136.76 | 93.47 | 137.58 | 1.37 | 10 | 10 | 20 | 15 | 8 | 23 |
| Tissue type/TBV | Total white matter/TBV | 23.63 | 24.59 | 23.99 | 24.38 | 25.22 | 1.77 | 7 | 7 | 14 | 10 | 5 | 15 |
|  | Total grey matter/TBV | 76.37 | 75.41 | 76.01 | 75.62 | 74.78 | 1.77 | 7 | 7 | 14 | 10 | 5 | 15 |
| Cortical regions  /TBV | R Occipital/TBV | 7.12 | 6.69 | 7.97 | 7.34 | 7.13 | 0.32 | 151 | 151 | 302 | 226 | 113 | 339 |
|  | R Temporal Auditory/TBV | 3.11 | 2.98 | 2.99 | 2.95 | 2.87 | 1.57 | 8 | 8 | 16 | 12 | 6 | 18 |
|  | R Subcortical/TBV | 3.24 | 3.50 | 3.57 | 3.16 | 3.39 | 0.13 | 947 | 947 | 1894 | 1421 | 711 | 2132 |
|  | R Frontal/TBV | 4.25 | 4.80 | 4.34 | 4.46 | 4.52 | 0.29 | 185 | 185 | 370 | 278 | 139 | 417 |
|  | R Cerebellum/TBV | 2.93 | 2.44 | 1.76 | 2.34 | 2.71 | 0.77 | 28 | 28 | 56 | 42 | 21 | 63 |
|  | R Insula/TBV | 0.50 | 0.60 | 0.52 | 0.48 | 0.44 | 1.43 | 9 | 9 | 18 | 14 | 7 | 21 |
|  | R Cingulate/TBV | 1.43 | 1.27 | 1.55 | 1.53 | 1.38 | 0.56 | 51 | 51 | 102 | 76 | 38 | 114 |
|  | R Parietal/TBV | 5.69 | 5.17 | 5.62 | 5.68 | 5.31 | 0.94 | 19 | 19 | 38 | 28 | 14 | 42 |
|  | R Prefrontal/TBV | 4.17 | 4.47 | 3.96 | 4.11 | 4.13 | 0.24 | 281 | 281 | 562 | 422 | 211 | 633 |
|  | R Corpus Callosum/TBV | 0.25 | 0.17 | 0.21 | 0.25 | 0.14 | 1.55 | 8 | 8 | 16 | 12 | 6 | 18 |
|  | R Temporal Visual/TBV | 4.10 | 4.03 | 4.01 | 4.39 | 4.48 | 1.58 | 8plos | 8 | 16 | 12 | 6 | 18 |
|  | R Temporal Limbic/TBV | 1.54 | 1.71 | 1.82 | 1.70 | 1.65 | 0.40 | 101 | 101 | 202 | 150 | 75 | 225 |
|  | R Pons and Medulla/TBV | 0.11 | 0.14 | 0.08 | 0.12 | 0.14 | 1.20 | 12 | 12 | 24 | 18 | 9 | 27 |
|  | L Occipital/TBV | 7.84 | 6.76 | 8.08 | 7.20 | 6.45 | 1.47 | 9 | 9 | 18 | 13 | 7 | 20 |
|  | L Temporal Auditory/TBV | 2.92 | 2.84 | 2.95 | 2.90 | 2.89 | 0.22 | 322 | 322 | 644 | 483 | 242 | 725 |
|  | L Subcortical/TBV | 3.13 | 3.30 | 3.30 | 2.95 | 2.96 | 1.19 | 13 | 13 | 26 | 19 | 10 | 29 |
|  | L Frontal/TBV | 3.85 | 4.52 | 4.23 | 4.22 | 4.36 | 0.63 | 41 | 41 | 82 | 61 | 31 | 92 |
|  | L Cerebellum/TBV | 2.93 | 2.44 | 1.93 | 2.42 | 2.64 | 0.56 | 51 | 51 | 102 | 76 | 38 | 114 |
|  | L Insula/TBV | 0.53 | 0.53 | 0.61 | 0.46 | 0.42 | 1.54 | 8 | 8 | 16 | 12 | 6 | 18 |
|  | L Cingulate/TBV | 1.48 | 1.28 | 1.41 | 1.41 | 1.14 | 1.88 | 6 | 6 | 12 | 9 | 5 | 14 |
|  | L Parietal/TBV | 5.18 | 5.22 | 5.42 | 5.22 | 5.08 | 1.44 | 9 | 9 | 18 | 14 | 7 | 21 |
|  | L Prefrontal/TBV | 3.72 | 4.45 | 3.76 | 3.89 | 3.84 | 0.40 | 97 | 97 | 194 | 146 | 73 | 219 |
|  | L Corpus Callosum/TBV | 0.14 | 0.11 | 0.12 | 0.09 | 0.12 | 0.43 | 85 | 85 | 170 | 128 | 64 | 192 |
|  | L Temporal Visual/TBV | 4.07 | 3.67 | 3.58 | 3.99 | 4.03 | 0.91 | 20 | 20 | 40 | 30 | 15 | 45 |
|  | L Temporal Limbic/TBV | 1.61 | 1.61 | 1.56 | 1.74 | 1.79 | 1.65 | 7 | 7 | 14 | 11 | 6 | 17 |
|  | L Pons and Medulla/TBV | 0.14 | 0.18 | 0.05 | 0.20 | 0.22 | 1.20 | 12 | 12 | 24 | 18 | 9 | 27 |
| Subcortical regions  /TBV | R Hippocampus/TBV | 0.40 | 0.47 | 0.58 | 0.45 | 0.43 | 0.58 | 48 | 48 | 96 | 72 | 36 | 108 |
|  | R Amygdala/TBV | 0.38 | 0.32 | 0.44 | 0.36 | 0.36 | 0.27 | 210 | 210 | 420 | 316 | 158 | 474 |
|  | R Caudate/TBV | 0.53 | 0.60 | 0.57 | 0.63 | 0.59 | 0.14 | 759 | 759 | 1518 | 1138 | 569 | 1707 |
|  | R Putamen/TBV | 1.02 | 1.08 | 1.11 | 0.92 | 1.06 | 0.33 | 144 | 144 | 288 | 215 | 108 | 323 |
|  | L Hippocampus/TBV | 0.41 | 0.43 | 0.37 | 0.45 | 0.45 | 1.17 | 13 | 13 | 26 | 19 | 10 | 29 |
|  | L Amygdala/TBV | 0.40 | 0.38 | 0.38 | 0.36 | 0.36 | 0.94 | 19 | 19 | 38 | 29 | 15 | 44 |
|  | L Caudate/TBV | 0.51 | 0.54 | 0.54 | 0.57 | 0.52 | 0.96 | 18 | 18 | 36 | 28 | 14 | 42 |
|  | L Putamen/TBV | 1.09 | 1.09 | 1.09 | 1.01 | 1.05 | 0.45 | 80 | 80 | 160 | 120 | 60 | 180 |
| Tissue type/ICV | Total white matter/ICV | 19.83 | 20.93 | 20.15 | 20.40 | 20.97 | 1.30 | 11 | 11 | 22 | 16 | 8 | 24 |
|  | Total grey matter/ICV | 64.09 | 64.17 | 63.83 | 63.28 | 62.19 | 2.02 | 5 | 5 | 10 | 8 | 4 | 12 |
| Cortical regions  /ICV | R Occipital/ICV | 5.98 | 5.70 | 6.69 | 6.14 | 5.93 | 0.53 | 58 | 58 | 116 | 86 | 43 | 129 |
|  | R Temporal Auditory/ICV | 2.61 | 2.54 | 2.51 | 2.47 | 2.39 | 1.74 | 7 | 7 | 14 | 10 | 5 | 15 |
|  | R Subcortical/ICV | 2.72 | 2.97 | 2.99 | 2.64 | 2.82 | 0.11 | 1402 | 1402 | 2804 | 2103 | 1052 | 3155 |
|  | R Frontal/ICV | 3.56 | 4.08 | 3.64 | 3.74 | 3.76 | 0.03 | 23207 | 23207 | 46414 | 34810 | 17405 | 52215 |
|  | R Cerebellum/ICV | 2.46 | 2.07 | 1.48 | 1.96 | 2.25 | 0.71 | 33 | 33 | 66 | 49 | 25 | 74 |
|  | R Insula/ICV | 0.42 | 0.51 | 0.44 | 0.40 | 0.37 | 1.42 | 9 | 9 | 18 | 14 | 7 | 21 |
|  | R Cingulate/ICV | 1.20 | 1.08 | 1.30 | 1.28 | 1.15 | 0.74 | 30 | 30 | 60 | 45 | 23 | 68 |
|  | R Parietal/ICV | 4.77 | 4.40 | 4.72 | 4.76 | 4.42 | 1.28 | 11 | 11 | 22 | 16 | 8 | 24 |
|  | R Prefrontal/ICV | 3.50 | 3.80 | 3.33 | 3.44 | 3.44 | 0.44 | 81 | 81 | 162 | 122 | 61 | 183 |
|  | R Corpus Callosum/ICV | 0.21 | 0.14 | 0.18 | 0.21 | 0.12 | 1.59 | 8 | 8 | 16 | 12 | 6 | 18 |
|  | R Temporal Visual/ICV | 3.44 | 3.43 | 3.37 | 3.68 | 3.72 | 1.53 | 8 | 8 | 16 | 12 | 6 | 18 |
|  | R Temporal Limbic/ICV | 1.29 | 1.46 | 1.52 | 1.42 | 1.37 | 0.57 | 49 | 49 | 98 | 74 | 37 | 111 |
|  | R Pons and Medulla/ICV | 0.09 | 0.12 | 0.06 | 0.10 | 0.12 | 1.13 | 14 | 14 | 28 | 20 | 10 | 30 |
|  | L Occipital/ICV | 6.58 | 5.75 | 6.78 | 6.03 | 5.36 | 1.58 | 8 | 8 | 16 | 12 | 6 | 18 |
|  | L Temporal Auditory/ICV | 2.45 | 2.41 | 2.48 | 2.43 | 2.40 | 1.27 | 11 | 11 | 22 | 17 | 9 | 26 |
|  | L Subcortical/ICV | 2.62 | 2.81 | 2.77 | 2.47 | 2.46 | 1.25 | 12 | 12 | 24 | 17 | 9 | 26 |
|  | L Frontal/ICV | 3.23 | 3.84 | 3.55 | 3.53 | 3.63 | 0.39 | 104 | 104 | 208 | 156 | 78 | 234 |
|  | L Cerebellum/ICV | 2.46 | 2.08 | 1.62 | 2.03 | 2.19 | 0.48 | 69 | 69 | 138 | 103 | 52 | 155 |
|  | L Insula/ICV | 0.44 | 0.45 | 0.51 | 0.39 | 0.35 | 1.57 | 8 | 8 | 16 | 12 | 6 | 18 |
|  | L Cingulate/ICV | 1.24 | 1.09 | 1.18 | 1.18 | 0.95 | 1.96 | 6 | 6 | 12 | 8 | 4 | 12 |
|  | L Parietal/ICV | 4.35 | 4.44 | 4.55 | 4.37 | 4.23 | 1.67 | 7 | 7 | 14 | 10 | 5 | 15 |
|  | L Prefrontal/ICV | 3.13 | 3.79 | 3.16 | 3.26 | 3.19 | 0.51 | 61 | 61 | 122 | 91 | 46 | 137 |
|  | L Corpus Callosum/ICV | 0.11 | 0.09 | 0.10 | 0.08 | 0.10 | 0.35 | 127 | 127 | 254 | 190 | 95 | 285 |
|  | L Temporal Visual/ICV | 3.41 | 3.12 | 3.00 | 3.34 | 3.36 | 0.77 | 28 | 28 | 56 | 42 | 21 | 63 |
|  | L Temporal Limbic/ICV | 1.35 | 1.37 | 1.31 | 1.46 | 1.49 | 1.59 | 8 | 8 | 16 | 12 | 6 | 18 |
|  | L Pons and Medulla/ICV | 0.12 | 0.16 | 0.04 | 0.16 | 0.19 | 1.16 | 13 | 13 | 26 | 20 | 10 | 30 |
| Subcortical regions  /ICV | R Hippocampus/ICV | 0.33 | 0.40 | 0.49 | 0.38 | 0.36 | 0.65 | 39 | 39 | 78 | 58 | 29 | 87 |
|  | R Amygdala/ICV | 0.32 | 0.27 | 0.37 | 0.30 | 0.30 | 0.38 | 108 | 108 | 216 | 162 | 81 | 243 |
|  | R Caudate/ICV | 0.44 | 0.51 | 0.48 | 0.53 | 0.49 | 0.04 | 8520 | 8520 | 17040 | 12780 | 6390 | 19170 |
|  | R Putamen/ICV | 0.86 | 0.92 | 0.93 | 0.77 | 0.88 | 0.15 | 726 | 726 | 1452 | 1088 | 544 | 1632 |
|  | L Hippocampus/ICV | 0.34 | 0.37 | 0.31 | 0.37 | 0.38 | 1.04 | 16 | 16 | 32 | 24 | 12 | 36 |
|  | L Amygdala/ICV | 0.34 | 0.33 | 0.32 | 0.30 | 0.30 | 1.09 | 15 | 15 | 30 | 22 | 11 | 33 |
|  | L Caudate/ICV | 0.43 | 0.46 | 0.46 | 0.48 | 0.43 | 1.18 | 13 | 13 | 26 | 19 | 10 | 29 |
|  | L Putamen/ICV | 0.91 | 0.93 | 0.92 | 0.85 | 0.88 | 0.70 | 33 | 33 | 66 | 50 | 25 | 75 |

Total brain volume (TBV). Intracranial volume (ICV). Cerebrospinal fluid (CSF). Control (Ctr). *These sample size estimates were performed for the observed effect sizes with 80% power at the two-sided 0.05 significance level
