## Supplementary material for "Quantitative definition of neurobehavior, vision, hearing and brain volumes in macaques congenitally exposed to Zika virus": S11 table

Supplementary Table 11. Fetus/Infant tissue viral loads.

| Organ System | Tissue Name | ZIKV-exposed fetus/infant tissue viral loads at necropsy | | | | |
| --- | --- | --- | --- | --- | --- | --- |
|  |  | 416597 | 424847 | 499874 | 527421 | 226691 |
| Digestive System | tongue | ND | ND | ND | ND | ND |
|  | esophagus | ND | ND | ND | ND | ND |
|  | stomach | ND | ND | ND | ND | ND |
|  | duodenum | ND | ND | ND | ND | ND |
|  | jejunum | ND | ND | ND | ND | ND |
|  | ileum | ND | ND | ND | ND | ND |
|  | colon | ND | ND | ND | ND | ND |
|  | cecum | ND | ND | ND | ND | ND |
|  | bile | NC | NC | ND | ND | ND |
|  | liver | ND | ND | ND | ND | ND |
| Renal | kidney | ND | ND | ND | ND | ND |
|  | urinary bladder | ND | ND | ND | ND | ND |
| Cardiovascular | heart | ND | ND | ND | ND | ND |
|  | pericardium | ND | ND | ND | ND | ND |
|  | aorta thoracic | ND | ND | ND | ND | ND |
| Connective | omentum | NC | ND | ND | ND | ND |
|  | epidermis/dermis | ND | ND | ND | ND | ND |
|  | bone marrow | ND | NC | ND | ND | ND |
| Immune | spleen | ND | ND | ND | ND | ND |
|  | thymus | ND | ND | ND | ND | ND |
|  | axillary LN | NC | ND | ND | ND | ND |
|  | inguinal LN | NC | ND | ND | ND | ND |
|  | mesenteric LN | ND | ND | ND | ND | ND |
|  | submandibular LN | NC | ND | ND | ND | ND |
|  | tracheobronchial LN | NC | ND | ND | ND | ND |
|  | oropharyngeal LN | ND | ND | ND | ND | NC |
|  | articular cartilage | ND | ND | ND | ND | ND |
| Musculoskeletal | femur | ND | ND | ND | ND | NC |
|  | muscle-quadriceps | ND | ND | ND | ND | ND |
|  | trabecular bone | NC | NC | NC | NC | NC |
|  | tendon | NC | NC | NC | NC | NC |
| Pulmonary | lung | ND | ND | ND | ND | ND |
| Reproductive | uterus | NA | NA | NA | NC | ND |
|  | ovary | NA | NA | NA | ND | ND |
|  | seminal vesicle | ND | ND | ND | NA | NA |
|  | testis | ND | ND | ND | NA | NA |
| Nervous | cerebrum 1* | ND | ND | ND | ND | ND |
|  | cerebrum 2* | ND | NC | NC | NC | NC |
|  | cerebrum 4* | ND | ND | ND | ND | ND |
|  | cerebrum 7* | ND | ND | ND | ND | ND |
|  | cerebrum 10* | ND | ND | ND | ND | ND |
|  | cerebellum 1* | NC | ND | NC | NC | NC |
|  | cerebellum 2* | NC | ND | ND | ND | ND |
|  | CSF | NC | ND | ND | ND | ND |
|  | dura mater | ND | ND | ND | ND | ND |
|  | cervical spinal cord | ND | ND | ND | ND | ND |
|  | thoracic spinal cord | ND | ND | ND | ND | ND |
|  | lumbar spinal cord | ND | ND | ND | ND | ND |
| Ocular | cornea | ND | ND | ND | ND | ND |
|  | optic nerve | NC | ND | ND | ND | ND |
|  | retina | ND | ND | ND | ND | ND |
|  | sclera | ND | ND | ND | ND | ND |
|  | eye-aqueous humor | NC | NC | ND | ND | ND |
| Endocrine | adrenal gland | ND | ND | ND | ND | ND |
|  | pancreas | ND | ND | ND | ND | ND |
|  | pituitary gland | ND | ND | ND | ND | ND |
|  | thyroid | ND | ND | ND | ND | ND |

Not detected (ND). Not collected (NC). Not applicable (NA). Lymph node (LN). * Ten slices of the fetal/infant cerebrum (~5 mm in thickness) were prepared in the coronal plane with section 1 located most anteriorly and section 10 located most posteriorly, and three slices of the cerebellum were prepared in the sagittal plane with section 1 located most laterally and section 3 most medially.
