## Supplementary material for "Quantitative definition of neurobehavior, vision, hearing and brain volumes in macaques congenitally exposed to Zika virus": S12 table

Supplementary Table 12. Infant/fetus morphometric measurements at necropsy.

|  |  | ZIKV-exposed | | | | | Control | Reference ranges from Scott JA et al. 2015.* |
| --- | --- | --- | --- | --- | --- | --- | --- | --- |
|  |  | 416597 | 424847 | 499874 | 527421 | 226691 | 020501 |  |
| Age at necropsy | Postnatal age (day of life) | NA | 8 | 7 | 8 | 8 | 8 | 1 week |
|  | Corrected gestational age | 133 | 163 | 162 | 162 | 163 | 165 | Undefined |
| Measurement | Femur length, right (cm) | 4.2 | 6.3 | ND | 4.9 | 5.7 | 5.4 | NA |
|  | Femur length, left (cm) | 4.1 | 6.3 | ND | 5 | 5.7 | 5.4 | NA |
|  | Brain weight (g) | 40.982 | 58.429 | ND | 55.87 | 53.716 | 54.28 | NA |
|  | Head circumference (cm) | 16 | 19.9 | ND | 19.2 | 19.1 | 19 | 21.0 (0.7 SD) female; 21.0 (0.5 SD) male |
|  | Biparietal diameter (cm) | 4.2 | 4.8 | ND | 5 | 5 | 4.9 | 5.3 (0.2 SD) female; 5.4 (0.2 SD) male |
|  | Crown to rump length (cm) | 17.5 | 21 | ND | 23 | 19 | 19.8 | 21.4 (0.8 SD) female; 21.5 (0.9 SD) male |
|  | Crown to tail length (cm) | 27 | 32.5 | ND | 29 | 30.5 | 30.4 | NA |
|  | Eye weight, right (g) | 1.124 | ND | ND | 1.553 | 1.552 | 1.71 | NA |
|  | Eye weight, left (g) | 0.982 | 1.1 | ND | 1.731 | 1.579 | 1.59 | NA |

Not applicable (NA). Not done (ND). Standard deviation (SD).

*No reference ranges exist for infants delivered at 155gd or 133gd. The closest reference ranges come from a publication describing morphometric measurements for infants born at term, presumably over 165gd (4). Our term infant had smaller head cirumferences than the reference ranges listed in this publication, but this may simply be due to age differences, because our control infant had a head circumference within range of our ZIKV-exposed infants.
