## Supplementary material for "Quantitative definition of neurobehavior, vision, hearing and brain volumes in macaques congenitally exposed to Zika virus": S13 table

Supplementary table 13. Histopathological description of the lung and middle ear in fetuses/infants.

| Tissue Source | Organ System | Tissue Name | ZIKV-exposed | | | | | Control |
| --- | --- | --- | --- | --- | --- | --- | --- | --- |
|  |  |  | Fetus: 416597 | Infant: 424847 | Infant: 499874 | Infant: 527421 | Infant: 226691 | Infant: 020501 |
|  | Pulmonary | Lung | Moderate autolysis, moderate diffuse atelectasis, moderate numbers of squamous cells within alveoli, and variable amounts of cellular debris, consistent with dystocia and fetal distress in utero. Bacteria were not identified. | No significant lesions | Moderate to marked, multifocal, lymphoplasmacytic and neutrophilic interstitial pneumonia with moderate, multifocal, neutrophilic and necrotizing interstitial bronchiolitis. Bacteria were not identified. | Mild diffuse perivascular edema surrounding large pulmonary arteries. | Severe diffuse bilateral neutrophilic bronchopneumonia with atelectasis. Bacteria were not identified. | Small numbers of alveolar macrophages throughout alveoli. Occasional macrophages exhibit erythrophagocytosis and there are rare plasma cells within alveoli as well. A rare syncytial cell (multinucleated) is noted. |
|  | Ear | Ear | Left ear: Minimal neutrophilic otitis media with intraluminal squamous cells. Bacteria were not identified. Right ear: Minimal neutrophilic otitis media with intraluminal squamous cells. | No significant histologic lesions bilaterally | Moderate to marked subacute neutrophilic otitis media. Bacteria were not identified. | Left ear: Minimal neutrophilic otitis media. Bacteria not identified. Right ear: no significant histologic lesions | No significant histologic lesions bilaterally | No significant histologic lesions bilaterally |

Not collected (NC)
