## Supplementary material for "Quantitative definition of neurobehavior, vision, hearing and brain volumes in macaques congenitally exposed to Zika virus": S14 table

Supplementary Table 14. Histopathological description of fetal/infant tissues excluding the lung and ear.

| Organ System | Tissue Name | ZIKV-exposed | | | | | Control |
| --- | --- | --- | --- | --- | --- | --- | --- |
|  |  | Fetus: 416597 | Infant: 424847 | Infant: 499874 | Infant: 527421 | Infant: 226691 | Infant: 020501 |
| Extraembryonic | amniotic membrane | No significant lesions | NC | Amniotic membranes are unremarkable. Chorionic membranes have small foci of lymphocytes scattered throughout the sections. | No significant lesions | NC | No significant lesions |
|  | umbilical cord | Mild focal suppurative vasculitis | NC | No significant lesions | No significant lesions | No significant lesions | Pseudothrombus (not adhered to the endothelium) in a single vessel, otherwise no significant lesions |
| Digestive System | tongue | Moderate to severe diffuse autolysis of tissue sections with no other significant lesions | No significant lesions | No significant lesions | No significant lesions | NC | No significant lesions |
|  | salivary gland | NC | NC | NC | No significant lesions | No significant lesions | No significant lesions |
|  | esophagus | Moderate to severe diffuse autolysis of tissue sections with no other significant lesions | No significant lesions | No significant lesions | No significant lesions | No significant lesions | No significant lesions |
|  | stomach | Moderate to severe diffuse autolysis of tissue sections with no other significant lesions | No significant lesions | No significant lesions | No significant lesions | Mild lymphocytic gastritis with nodular lymphoid hyperplasia. | No significant lesions |
|  | duodenum | NC | No significant lesions | No significant lesions | No significant lesions | No significant lesions | No significant lesions |
|  | jejunum | Moderate to severe diffuse autolysis of tissue sections with no other significant lesions | No significant lesions | No significant lesions | No significant lesions | No significant lesions | No significant lesions |
|  | ileum | NC | No significant lesions | No significant lesions | NC | No significant lesions | No significant lesions |
|  | colon | Moderate to severe diffuse autolysis of tissue sections with no other significant lesions | No significant lesions | No significant lesions | No significant lesions | No significant lesions | No significant lesions |
|  | cecum | Moderate to severe diffuse autolysis of tissue sections with no other significant lesions |  | No significant lesions | No significant lesions | No significant lesions | No significant lesions |
|  | liver | NC | Minimal lymphoplasmacytic centrilobular and periportal hepatitis. | Lymphoplasmacytic perivasculitis. Moderate, multifocal, portal and perivascular lymphoplasmacytic and neutrophilic hepatitis. | No significant lesions | No significant lesions | Minimal multifocal neutrophilic hepatitis |
|  | gall bladder | NC | NC | NC | No significant lesions | NC | No significant lesions |
| Renal | kidney | Moderate to severe diffuse autolysis of tissue sections with no other significant lesions | No significant lesions | Bilateral, focal, renal cysts. | No significant lesions | No significant lesions | No significant lesions |
|  | urinary bladder | Moderate to severe diffuse autolysis of tissue sections with no other significant lesions | No significant lesions | No significant lesions | No significant lesions | No significant lesions | No significant lesions |
| Cardiovascular | heart | Moderate to severe diffuse autolysis of tissue sections with no other significant lesions | No significant lesions | No significant lesions | No significant lesions | No significant lesions | No significant lesions |
|  | pulmonary artery | NC | No significant lesions | NC | No significant lesions | NC | NC |
|  | mesenteric arteries | NC | No significant lesions | NC | NC | NC | NC |
|  | pericardium | Moderate to severe diffuse autolysis of tissue sections with no other significant lesions | No significant lesions | Lymphoplasmacytic perivasculitis Minimal, perivascular and random lymphoplasmacytic, neutrophilic pericarditis. | No significant lesions | No significant lesions | Small aggregates of mononuclear cells within capillaries |
|  | aorta | Moderate to severe diffuse autolysis of tissue sections with no other significant lesions | No significant lesions | Mild to moderate multifocal, lymphoplasmacytic and neutrophilic adventitia-vascular periarteritis and vasa-vasorum vasculitis. | No significant lesions | No significant lesions | No significant lesions |
| Connective | omentum | NC | No significant lesions | No significant lesions | No significant lesions | No significant lesions | No significant lesions |
|  | brown fat | Moderate to severe diffuse autolysis of tissue sections with no other significant lesions | No significant lesions | NC | No significant lesions | No significant lesions | NC |
|  | skin | Moderate to severe diffuse autolysis of tissue sections with no other significant lesions | No significant lesions | No significant lesions | No significant lesions | No significant lesions | No significant lesions |
| Immune | bone marrow | Moderate to severe diffuse autolysis of tissue sections with no other significant lesions | Minimal to mild inflammation and erythroid hyperplasia, and mild granulocytic hyperplasia. | Immature and mature polymorphonuclear cells with small islands of erythropoiesis. | NC | No significant lesions | NC |
|  | axillary LN | NC | No significant lesions | No significant lesions | No significant lesions | No significant lesions | Mild diffuse neutrophilic lymphadenitis |
|  | inguinal LN | NC | No significant lesions | No significant lesions | No significant lesions | No significant lesions | Mild diffuse neutrophilic lymphadenitis |
|  | lymph node, nos | Moderate to severe diffuse autolysis of tissue sections with no other significant lesions | NC | NC | NC | NC | NC |
|  | mesenteric LN | NC | Minimal neutrophilic lymphadenitis with hemosiderosis | No significant lesions | No significant lesions | No significant lesions | Minimal neutrophilic inflammation with hemosiderosis |
|  | pancreatic LN | NC | No significant lesions | NC | No significant lesions | No significant lesions | Moderate diffuse neutrophilic lymphadenitis |
|  | parotid LN | NC | No significant lesions | NC | NC | NC | NC |
|  | splenic LN | NC | NC | NC | No significant lesions | NC | Moderate diffuse neutrophilic lymphadenitis |
|  | submandibular LN | NC | No significant lesions | No significant lesions | No significant lesions | NC | Mild diffuse neutrophilic lymphadenitis |
|  | thoracic LN | NC | No significant lesions | NC | NC | NC | NC |
|  | tonsil/oropharyngeal LN | NC | No significant lesions | No significant lesions | NC | No significant lesions | NC |
|  | tracheobronchial LN | Moderate to severe diffuse autolysis with no other significant lesions. | No significant lesions | Moderate to marked neutrophilic lymphadenitis | NC | Minimal neutrophilic lymphadenitis. | Mild neutrophilic lymphadenitis |
|  | spleen | NC | No significant lesions | Mild increase of neutrophils in the red pulp | Mild lymphoid hyperplasia of periarterial lymphatic sheaths | Moderate diffuse neutrophilic splenitis | No significant lesions |
|  | thymus | Moderate to severe diffuse autolysis of tissue sections with no other significant lesions | No significant lesions | No significant lesions | No significant lesions | Minimal multifocal suppurative thymitis | No significant lesions |
| Musculoskeletal | articular cartilage | Moderate to severe diffuse autolysis of tissue sections with no other significant lesions | No significant lesions | No significant lesions | No significant lesions | No significant lesions | No significant lesions |
|  | femur | NC | No significant lesions | No significant lesions | No significant lesions | No significant lesions | NC |
|  | ligament of the head of the femur | NC | No significant lesions | NC | NC | No significant lesions | NC |
|  | quadriceps muscle | Moderate to severe diffuse autolysis of tissue sections with no other significant lesions | No significant lesions | No significant lesions | No significant lesions | No significant lesions | No significant lesions |
|  | trabecular bone | NC | No significant lesions | NC | NC | No significant lesions | NC |
|  | tendon | NC | NC | NC | NC | No significant lesions | NC |
| Respiratory | larynx | NC | NC | Mild-moderate, neutrophilic laryngitis | NC | NC | NC |
|  | oropharynx | NC | NC | Moderate multifocal neutrophilic and lymphoplasmacytic oropharyngitis | NC | No significant lesions | No significant lesions |
|  | trachea | NC | No significant lesions | No significant lesions | NC | NC | NC |
| Reproductive | cervix | NA | NA | NA | NC | No significant lesions | No significant lesions |
|  | uterus | NA | NA | NA | No significant lesions | No significant lesions | No significant lesions |
|  | vagina | NA | NA | NA | NC | No significant lesions | No significant lesions |
|  | fallopian tube | NA | NA | NA | No significant lesions | No significant lesions | No significant lesions |
|  | fimbria | NA | NA | NA | NC | No significant lesions | No significant lesions |
|  | ovary | NA | NA | NA | No significant lesions | No significant lesions | No significant lesions |
|  | epididymis | NC | No significant lesions | Lymphoplasmacytic perivasculitis | NA | NA | NA |
|  | prostate | NC | No significant lesions | NC | NA | NA | NA |
|  | seminal vesicle | Moderate to severe diffuse autolysis of tissue sections with no other significant lesions | No significant lesions | No significant lesions | NA | NA | NA |
|  | testis | Moderate to severe diffuse autolysis of tissue sections with no other significant lesions | No significant lesions | Multiple blood vessels with small perivascular aggregates of lymphocytes plasma cells and neutrophils. | NA | NA | NA |
| Nervous | cerebrum | Moderate to severe diffuse autolysis of tissue sections with no other significant lesions. Herniation of the cerebellum with compression of the spinal cord without hemorrhage or neuronal necrosis is consistent with a post mortem event. | No significant lesions | No significant lesions | No significant lesions | No significant lesions | No significant lesions |
|  | cerebellum | Moderate to severe diffuse autolysis of tissue sections with no other significant lesions | No significant lesions | NC | No significant lesions | No significant lesions | No significant lesions |
|  | dura mater | Moderate to severe diffuse autolysis of tissue sections with no other significant lesions | No significant lesions | No significant lesions | No significant lesions | NC | NC |
|  | cervical spinal cord | Moderately autolyzed with increased numbers of mononuclear cells throughout the parenchyma | No significant lesions | No significant lesions | No significant lesions | No significant lesions | No significant lesions |
|  | thoracic spinal cord | Moderately autolyzed with increased numbers of mononuclear cells throughout the parenchyma |  | No significant lesions | No significant lesions | No significant lesions | No significant lesions |
|  | lumbar spinal cord | Moderately autolyzed with increased numbers of mononuclear cells throughout the parenchyma | No significant lesions | No significant lesions | No significant lesions | No significant lesions | No significant lesions |
|  | cauda equina | NC | NC | No significant lesions | NC | NC | NC |
|  | brachial nerve | NC | No significant lesions | NC | NC | NC | NC |
|  | mesenteric ganglion | NC | NC | NC | NC | No significant lesions | NC |
| Ocular | globe | Globe is markedly distorted, but presents normal overall size and shape. Histologically the tissues are severely autolyzed, precluding adequate microscopic analysis. | Normal overall size and shape. Histologically no developmental or inflammatory lesions. | Normal overall size and shape. Histologically no developmental or inflammatory lesions. | Normal overall size and shape. Histologically no developmental or inflammatory lesions. | Normal overall size and shape. Histologically no developmental or inflammatory lesions. | Normal overall size and shape. Histologically no developmental or inflammatory lesions. |
| Endocrine | adrenal gland | Moderate to severe diffuse autolysis of tissue sections with no other significant lesions | No significant lesions | No significant lesions | No significant lesions | No significant lesions | No significant lesions |
|  | pancreas | NC | No significant lesions | No significant lesions | No significant lesions | No significant lesions | No significant lesions |
|  | pituitary gland | NC | No significant lesions | No significant lesions | No significant lesions | No significant lesions | No significant lesions |
|  | thyroid | Moderate to severe diffuse autolysis of tissue sections with no other significant lesions | No significant lesions | No significant lesions | No significant lesions | No significant lesions | No significant lesions |

Not collected (NC). Not applicable (NA). Lymph node (LN).
