## Supplementary material for "Quantitative definition of neurobehavior, vision, hearing and brain volumes in macaques congenitally exposed to Zika virus": S15 table

Supplementary Table 15. Summary of sample size estimates for all quantitative infant exams

| Infant Exam | Number of quantitative exam parameters that meet sample size allocations smaller than 8 or 14 infants per group* | | Figure/Table with observed effect & sample sizes |
| --- | --- | --- | --- |
|  | 8 infants per group (effect size >1.5) | 14 infants per group (effect size >1.1) |  |
| Feeding volume and weight gain | 0/2 | 1/2 | Figure 6 |
| Optical coherence tomography | 5/10 | 5/10 | S8 table |
| Visual electrophysiology | 2/18 | 8/18 | S9 table |
| Auditory brainstem response testing | 0/6 | 6/6 | Figure 10 |
| Volumetric brain analysis^#^ | 10/40 | 19/40 | S10 table |
| All exams combined | 17/76 (22%) | 39/76 (51%) | - |

*The denominator is the total number of parameters measured for a specific exam. The numerator is the number of parameters with an observed effect size greater than the noted effect size for the column.
#Volumetric brain analysis includes the following parameters described in table S10: total brain volumes and region volumes corrected for TBV.
